# Effect of contact pressure on porcine postmortem brain tissue impedance

**DOI:** 10.1101/2022.08.04.502741

**Authors:** Lucas Poßner, Florian Wilhelmy, Uwe Pliquett, Thomas R. Knösche, Konstantin Weise, Dirk Lindner

## Abstract

In this experimental study we demonstrate the influence of contact pressure on porcine postmortem brain tissue impedance using a movable electrode array and a load cell. We show that the variation of the contact pressure between the tissue and the measurement probe leads to a coefficient of variation in the measured impedance of under 3%. Its influence can therefore be neglected. Further, we fit the measured impedances to an equivalent circuit model and compare the resistance as well as the cell density of grey and white matter brain tissue based on the model parameters.

## I. Introduction

The knowledge of the conductivity and the permittivity of brain tissue in living human subjects is relevant for a variety of applications such as electroencephalography, magnetoencephalography, and brain stimulation techniques, which are routinely used in clinical and scientific practice. Moreover, the knowledge of these properties may be of use for the identification of pathological tissue, such as brain tumors. Existing studies in which these properties were investigated exhibit a variety of drawbacks such as an inappropriate measurement chain, the marginalization of important influencing variables or an insufficient number of subjects [1]. The electric properties of biological tissue can be measured using different techniques. Among the most direct and accurate is the measurement of the electric impedance using current application with a probe directly applied to the tissue and thus deriving the conductivity and permittivity from the voltage dropping across the object. We plan to conduct a study in which we acquire the impedance of the different tissues of the living human brain, including pathological tissue, during the resection of brain tumors using this technique. In former publications, we addressed some of the substantial challenges in the measurement of the impedance of brain tissue [2,3]. We provided a proof-of-principle study in which we distinguished grey and white matter of in-situ porcine brain tissue using impedance measurements [2] and proposed a novel measurement probe design to conduct impedance measurements using a four-electrode-configuration [3]. To facilitate intraoperative measurements, other influencing factors have to be considered: As the measurement probe has to be manually applied intraoperatively onto flexible, living tissue, the contact pressure between the probe and the tissue might vary considerably. This might influence the measured impedance significantly [4]. Here, we investigate this issue. Further, we fit the measured impedances to an equivalent circuit model and compare the resistance as well as the cell density of grey and white matter brain tissue based on the model parameters. Again, we use porcine postmortem brain tissue since it is widely available and its structure is comparable to human brain tissue [5].

## II. Methods

### A. Experimental setup of contact pressure influence evaluation

To evaluate the effect of contact pressure on grey and white matter brain impedance, we used a linear gold electrode array (Sciospec Scientific Instruments GmbH, Bennewitz, Germany) with an electrode length of 3.2 mm, an electrode width of 0.2 mm and an electrode distance of 0.6 mm (Fig. 1a) that was connected to an ISX-3 mini impedance analyzer (Sciospec Scientific Instruments GmbH, Bennewitz, Germany). The tissue was placed in a movable vessel with a diameter of 5 mm and the array was mounted at a fixed position above it (Fig. 1b and 1c). The contact pressure was varied using a stepper motor that is attached to a shaft that moves the vessel perpendicularly to the array. This resembles the conditions during a surgery where the surgent applies the measurement probe to the brain tissue of the subject. We measured the force on the tissue using a load cell that was connected to the vessel using a second shaft (Fig. 1d). We conducted measurements of grey and white matter porcine postmortem brain impedance using a four-point-configuration in a frequency range between 1 kHz and 1 MHz. The two current-carrying electrodes working (W) and counter (C) as well as the two voltage-pickup electrodes sensing (WS) and reference (R) were chosen to have the highest possible distance. This results in a distance of 3.6 mm between the current-carrying electrodes and a distance of 2.4 mm between the voltage-pickup electrodes (Fig. 1a). A current of 10 μA was applied. We recorded the average of 10 spectra in each measurement.

**Fig. 1.**
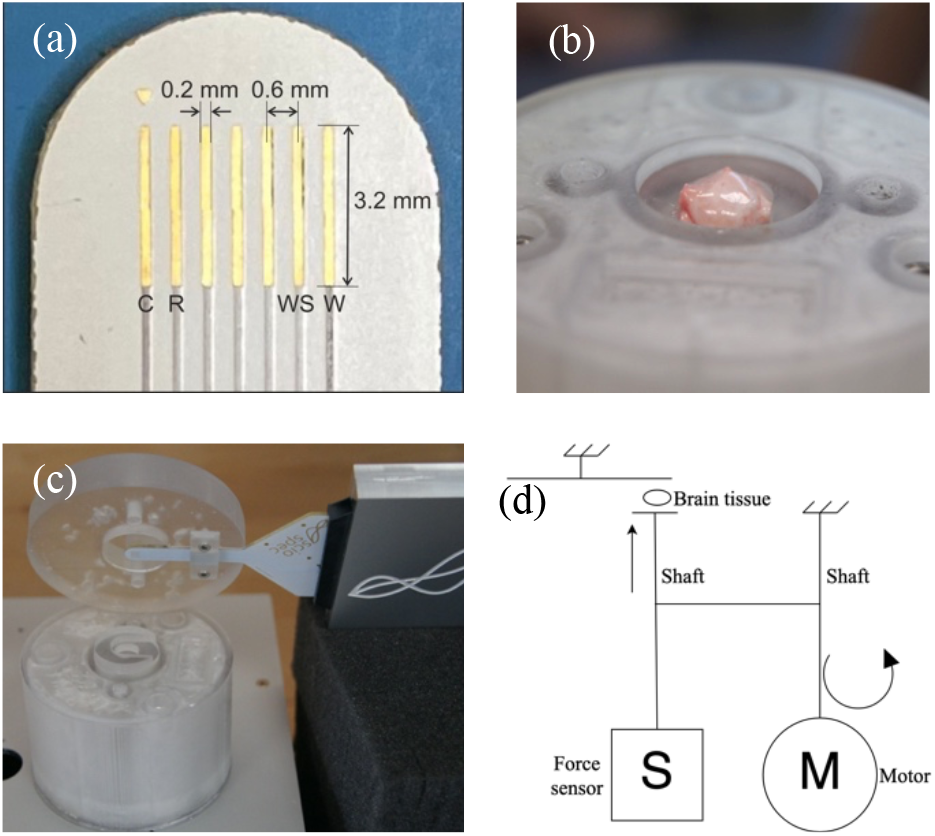
(a) Linear gold electrode array with electrode length of 3.2 mm, an electrode width of 0.2 mm and an electrode distance of 0.6 mm. (b) Movable vessel with porcine brain tissue placed inside. (c) Schematic of the measurement setup with the stepper motor (M) and the force sensor (S). (d) Linear gold electrode array with the working-(W) and counter-electrode (C) as well as the sensing-(WS) and reference-electrode (R).

### B. Evaluation of the contact pressure influence

We increased the pressure on the tissue by rotating the stepper motor in 90° steps (corresponding to 0.5 mm lift-off). The force offset was calibrated by measuring the output value of the load cell when the tissue had no contact to the electrode array. Additionally, we normalized the force to the output value when the tissue was maximally compressed and squeezed. It is worthwhile to note that such high forces lead to tissue damage and will never be exerted to living human brain tissue during surgery. After each step we measured the impedance. Since the contact area between the electrode array and the brain tissue stayed constant, the normalized force can be interpreted as the relative contact pressure that is applied to the tissue, which is expressed as *p*/*p*_0_. We fitted a linear function on the absolute value and the phase angle of the measured impedances with respect to the relative contact pressure at each frequency point. This can be expressed as:

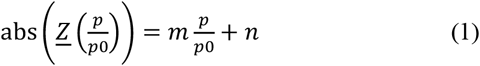

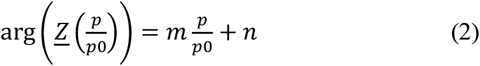

The relative contact pressure that is applied during intraoperative measurements can be modeled as a random variable *P*. As a probability density function for *P* we chose the Beta distribution, since both the relative contact pressure and the Beta distribution are defined on an interval *P* = *p*/*p*0 ∈ [0,1]. As parameters we chose *α* = 4 and *β* = 10 which yields to a right-skewed distribution of *P*, since we expect the applied relative contact pressure to be much more likely in the lower interval range. The random variable *P* has a mean of *μ*_*p*_ = 0.29 *p*/*p*_0_ and a standard deviation of *σ*_*p*_ = 0.12 *p*/*p*_0_ (Fig. 2).

**Fig. 2.**
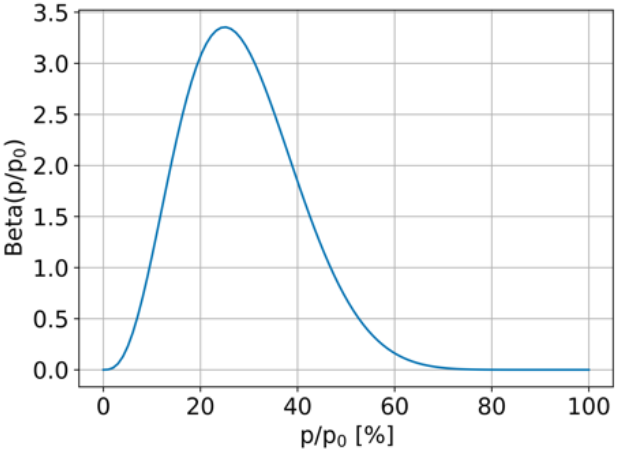
Probability density distribution of a random variable *P* that models the relative contact pressure 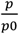. The distribution is a Beta distribution with parameters *α* = 4 and *β* = 10, a mean of *μ* = 0.29 *p*/*p*_0_ and a standard deviation of *σ* = 0.12 *p*/*p*_0_.

Since (1) and (2) depend on *P* = *p*/*p*0, the absolute value and the phase angle of the measured impedance at each frequency point are random variables. To derive a measure for the sensitivity of the measured impedance to the applied contact pressure, we sampled (1) and (2) for *n* = 1000 values of the relative contact pressure *p*/*p*0 and calculated the mean *μ*_abs,angle_ and the standard deviation *σ*_abs,angle_ of the resulting impedance. Additionally, we calculated the coefficient of variation *σ*/|*μ*|. In theory, the mean *μ*_abs,angle_ can be calculated as *mμ*_*p*_ + *n*, since we observed that the relationship between the absolute value, respectively phase angle, and the applied contact pressure is linear. Similarly, it can be shown that *σ*_abs,angle_ can be calculated as *mσ*_*p*_. Therefore, the mean *μ*_*p*_ and the variance *σ*_*p*_ are scaled by the slope *m* and the offset *n*.

### C. Equivalent circuit model

The measured impedances of grey and white matter tissue were fitted to an equivalent circuit model which consist of the parallel combination of a resistor *R*_1_ with a constant phase element *Q*_1_ and a single resistor *R*_2_ in series. The *R*_1_*Q*_1_ element can be used to model a dispersion with distributed time constants. The equivalent circuit model was chosen since it is a standard tissue model and it allows for an easy estimation of initial parameters that are used as start parameters in the Levenberg-Marquardt fitting algorithm. The equivalent circuit is shown in Fig. 3.

**Fig. 3.**
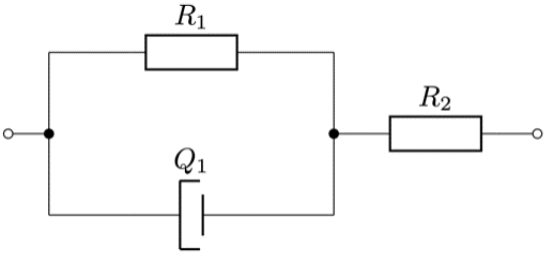
Equivalent circuit model that was used to fit the measured impedances of grey and white matter tissue. It consists of the parallel combination of a resistor *R*_1_ with a constant phase element *Q*_1_ and a single resistor *R*_2_ in series.

The impedance of a constant phase element is defined as:

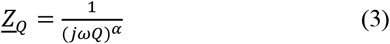

Therefore, the impedance of the equivalend circuit model is given by:

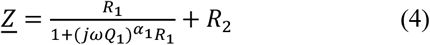

where *α*_1_ determines the broadness of the distribution of time constants. For *α* → 0 the distribution becomes infinetely broad and the circuit more resitive, for *α* → 1 the distribution becomes infinetely narrow and the circuit is more capacitive. For ω → 0 the impedance of the constant phase element goes to infinity. Therefore, the impedance of the equivalent circuit model at low frequencies becomes:

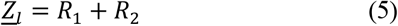

For ω → ∞ the impedance of the constant phase element goes to zero. Therefore, the impedance of the equivalent circuit model at high frequencies becomes:

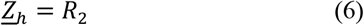

### D. Cell membrane density model

In [6] it was shown that an equivalent circuit model can be used to derive a measure reflecting the volume fraction of cells surrounded by insulating cell membranes in the tissue. The model consists of a parallel combination of a resistor (*R*_*ex*_) that models the extracellular conductive path and a capacitor (*C*_*m*_) that models the capacity of the cell membranes in series to a resistor (*R*_*in*_) that models the intracellular conductive path in the tissue. The equivalent circuit model is shown in Fig. 4. This model has a different topology than the equivalent circuit model that is used to fit the measured impedances. Even though the topology is different, the two models produce the same qualitative results and a unique mapping between their parameters exists.

**Fig. 4.**
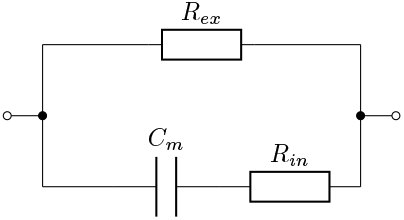
Equivalent circuit model that can be used to derive a measure that is a direct indicator for the volume fraction of cells surrounded by insulating cell membranes in the tissue. It consists of a parallel combination of a resistor (*R*_*ex*_) and a capacitor (*C*_*m*_) that in in series to a resistor (*R*_*in*_).

The impedance of the equivalent circuit model can be written as:

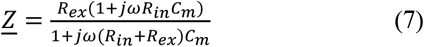

At low frequencies its impedance is only determined by the extracellular conductive path:

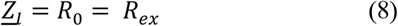

Respectivly, at high frequencies its impedance is determined by the extracellular and intracellualar conductive path:

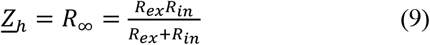

The parameters *R*_0_ and *R*_∞_ are used to define the measure

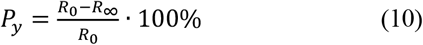

that is a direct indicator for the volume fraction of cells surrounded by insulating cell membranes in the tissue. The model that is based on (7) can be adapted to the model (4) that was used to fit the measured grey and white matter brain tissue impedances. Comparing (5) and (8), respectivly (6) and (9), yields a definition for *P*_*y*_ that is adapted to model (4):

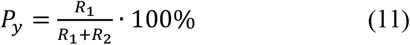

## III. Results

### A. Contact pressure influence

The absolute values as well as the phase angles of the measured impedances of grey and white matter brain tissue at different relative contact pressures are shown in Fig. 5.

**Fig. 5.**
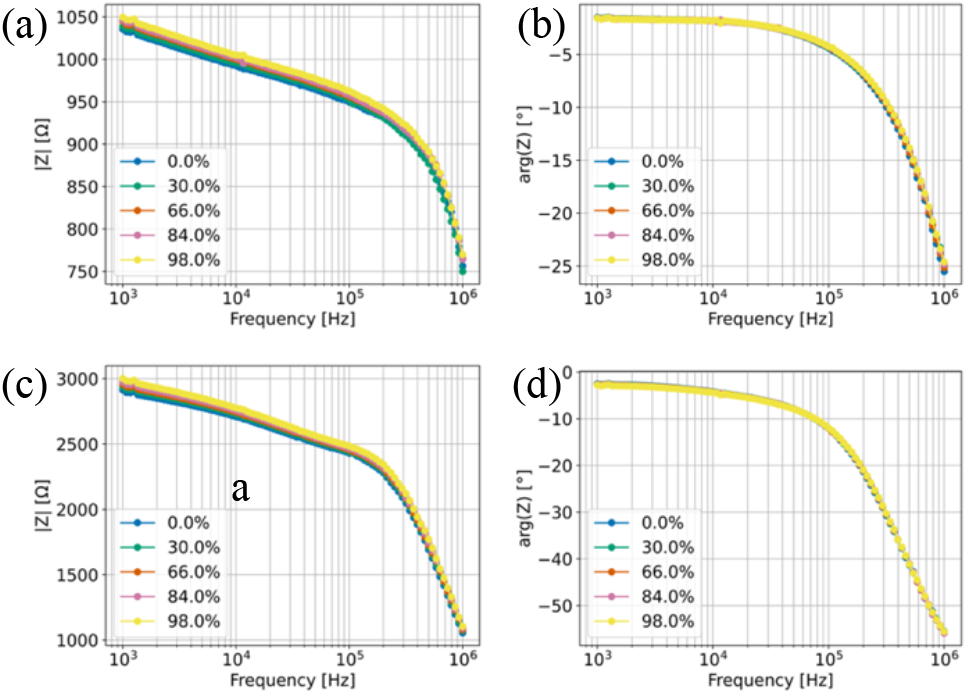
(a) Absolute value of the measured impedance of grey matter at different relative contact pressures. (b) Phase angle of the measured impedance of grey matter at different relative contact pressures. (c) Absolute value of the measured impedance of white matter at different relative contact pressures. (d) Phase angle of the measured impedance of white matter at different relative contact pressures.

The relative span of the absolute value and the phase angle with respect to the applied relative contact pressure is highest at 857 kHz. The measured impedance values and a linear fit at this frequency are shown in Fig. 6. The measured impedance at 100% relative contact pressure differs significantly from the rest of the measurement and was excluded in the further analysis.

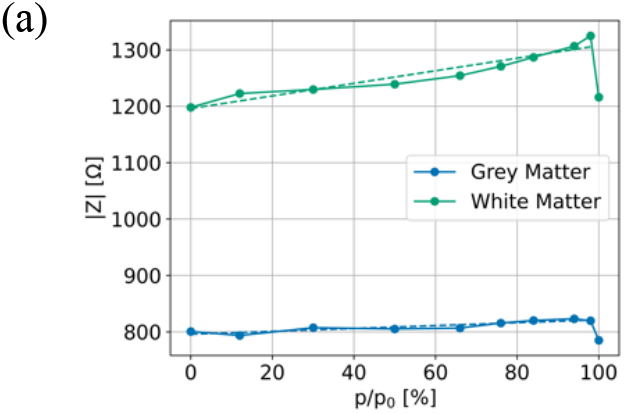
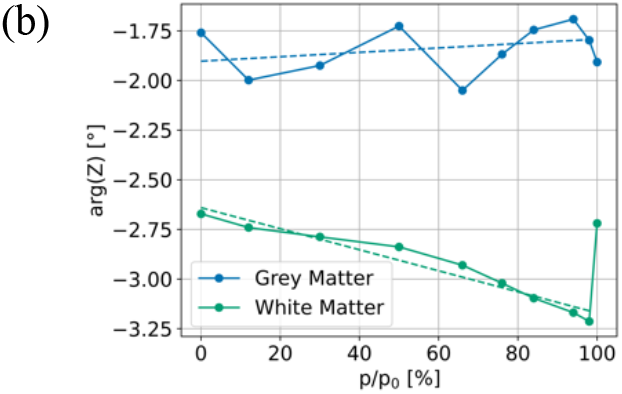

**Fig. 6.**
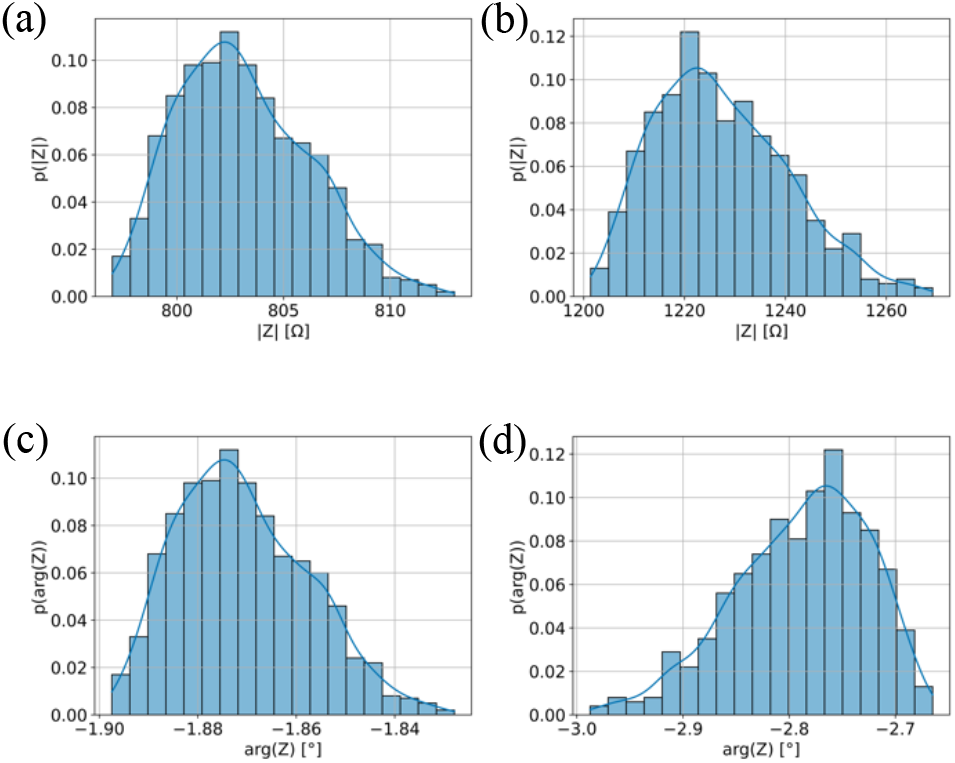
Simulation results obtained by sampling a linear function that was obtained from a fit to the absolute value and the phase angle of the measured impedances of grey and white matter at 857 kHz (a) Histogram for the absolute value of grey matter (b) Histogram for the absolute value of white matter (c) Histogram for the phase angle of grey matter (d) Histogram for the phase angle of grey matter

The linear functions that were obtained from the fit at 857 kHz were sampled with *n* = 1000 using the random variable *P* = *p*/*p*0. The histograms of the simulated values are shown in Fig. 7. The ordinate is density normalized such that the bins of the histrogram sum up to 1 i.e. it represents the probability of the abscissa values within a bin.

**Fig. 7.**
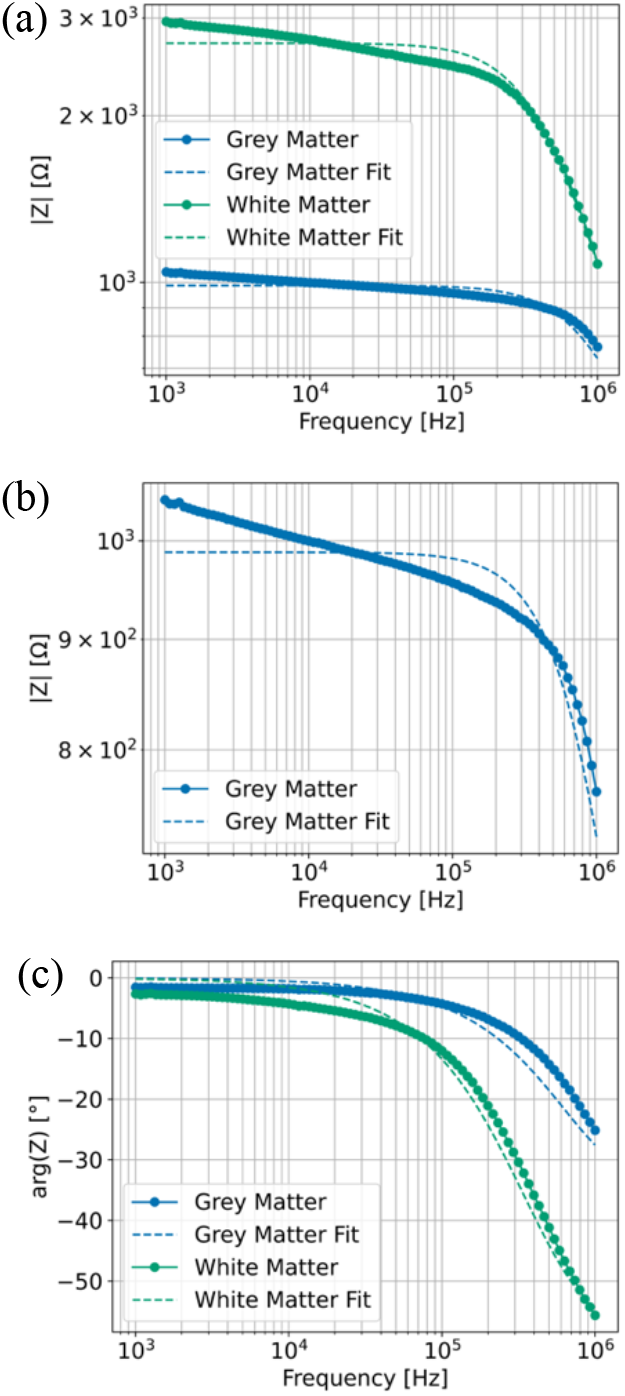
(a) Absolute values of the measured grey and white matter tissue impedances and equivalent circuit model fits. (b) Absolute value of the measured grey matter tissue impedance and equivalent circuit model fit. (c) Phase angles of the measured grey and white matter tissue impedances and equivalent circuit model fits.

The mean and the standard deviation as well as the coefficient of variation of the simulated values were calculated. Additionally, we calculated the theoretical mean and standard deviation by scaling the mean and standard deviation of the random variable *P* = *p*/*p*0 by the slope *m* and the offset *n* of the linear functions obtained from the fit. The simulated values are shown in Table I, Table II and Table III with the theoretical values in brackets.

**TABLE I.**
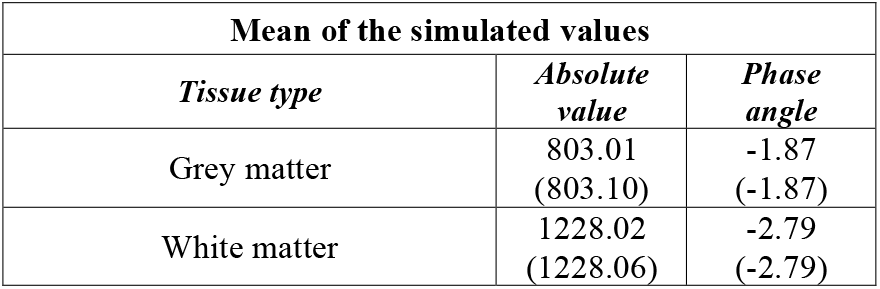
MEAN OF THE SIMULATED VALUES

| Mean of the simulated values |  |  |
| --- | --- | --- |
| Tissue type | Absolute value | Phase angle |
| Grey matter | 803.01<br>(803.10) | -1.87<br>(-1.87) |
| White matter | 1228.02<br>(1228.06) | -2.79<br>(-2.79) |

**TABLE II.** STANDARD DEVIATION OF THE SIMULATED VALUES

| Standard deviation of the simulated values |  |  |
| --- | --- | --- |
| Tissue type | Absolute value | Phase angle |
| Grey matter | 3.07<br>(2.98) | 0.013<br>(0.013) |
| White matter | 12.59<br>(13.00) | 0.060<br>(0.062) |

**TABLE III.** COEFFICIENT OF VARIATION OF THE SIMULATED VALUES

| Coefficient of variation of the simulated values |  |  |
| --- | --- | --- |
| Tissue type | Absolute value | Phase angle |
| Grey matter | 0.38 %<br>(0.37%) | 0.7%<br>(0.7%) |
| White matter | 1.02%<br>(1.05%) | 2.15%<br>(2.22%) |

### B. Equivalent circuit model fit

The measured impedances of grey and white matter tissue were fit to the equivalent circuit model (4) that is shown in Fig. 3. The measured impedances and the corresponding equivalent circuit model fits are shown in Fig. 8. The absolute values are shown in Fig. 8a, an enlarged plot of the absolute value of grey matter brain tissue is shown in Fig. 8b and the phase angels are shown in Fig. 8c. To evaluate the goodness-of-fit, the root-mean-square-deviation (RMSD) between the equivalent circuit model fit and the measured impedance was calculated. It is shown in Table III. The parameters of the equivalent circuit model fits are shown in Table IV.

**TABLE IV.**
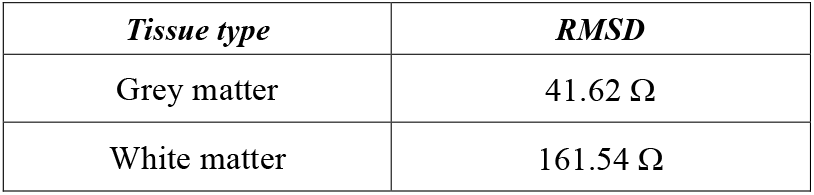
ROOT-MEAN-SQUARE-DEVIATION BETWEEN THE MEASURED IMPEDANCES THE EQUIVALENT CIRCUIT MODEL FITS

**TABLE V.** EQUIVALENT CIRCUIT MODEL PARAMETERS

| *Tissue type* | $R_1$<br>[$\Omega$] | $Q_1$<br>[$F^{-\alpha_1}$] | $\alpha_1$ | $R_2$<br>[$\Omega$] |
| --- | --- | --- | --- | --- |
| Grey matter | 675 | 8.55 e-10 | 0.92 | 313 |
| White matter | 2445 | 3.41 e-10 | 0.95 | 257 |

### C. Cell membrane density

The value of *P*_*y*_ was calculated from the parameters of the equivalent circuit model fit for grey and white matter brain tissue. For grey matter we found that *P*_*y*_ = 68.3% and for white matter we found that *P*_*y*_ = 90.4%.

## IV. Discussion

### A. Contact pressure influence

For both tissue types, the dependency of the measured impedance on the contact pressure is neglectable. The influence of the applied relative contact pressure is much smaller in comparison to the difference between grey and white matter. The standard deviation of the absolute value and the phase angle is small in comparison to its respective mean which leads to a coefficient of variation below 3%. The only significant change occurs when the tissue is squeezed and therefore damaged. This can be observed in Fig. 6. It is noted that pressures that lead to tissue damage will never be applied to living human brain tissue during surgery. From the results it can be concluded that the influence of the contact pressure on impedance measurements can be neglected.

### B. Equivalent circuit model fit

The root-mean-square-deviation between the measured impedance and the equivalent circuit model fit shows that the model that we used cannot fully describe the measured impedances of grey and white matter tissue. The measured impedances show a frequency dependence in the lower frequency range that is not matched by the equivalent circuit model that we used since it only accounts for a single dispersion. This could be due to parasitic elements in the measurement chain that were not fully compensated or certain effects that occur within the tissue or at the electrode-tissue-interface. The resistances *R*_1_ and *R*_2_ are larger for white matter brain tissue. This is plausible since grey matter is expected to have a higher conductivity than white matter. The other equivalent circuit model parameters do not show a significant difference.

### C. Cell membrane density

The value of *P*_*y*_ differs significantly between grey and white matter brain tissue. This gives evidence that the volume fraction of insulating cell membranes is higher in white matter than in grey matter. This could be due to the higher myelinization of white matter which leads to a higher density of insulating cells in the tissue. Since the normalized root mean square deviation of the equivalent circuit fit shows that the equivalent circuit model that we used cannot fully describe the measured impedances, the calculated values of *P*_*y*_ do have an increased uncertainty.

## V. Conclusion

It was shown that the influence of the contact pressure on impedance measurements is very small and can be neglected. The standard deviation of the absolute value and the phase angle due to contact pressure changes during impedance measurements is small in comparison to the mean which leads to a coefficient of variation below 3%. Since the mechanical structure of porcine brain tissue is comparable to human brain tissue, we hold our results as directly applicable to the practical use in a measurement during surgery. There are no significant changes in the measured impedance within the contact pressure range of a supposedly hand-held measurement probe, as long as electrode-tissue contact is complete i.e., the tissue covers the whole of the probe’s surface.

The equivalent circuit model that we used cannot fully describe the measured brain tissue impedances. Nevertheless, it meets the expectation that grey matter does have a higher conductivity than white matter.

Further, the equivalent circuit model was used to calculate a measure for the cell membrane density in the tissue. There is evidence that white matter does have a higher cell membrane density than grey matter. Since this deduction is based on a parameter with an increased uncertainty, it should be investigated further.

## Notes

### Competing Interest Statement

The authors have declared no competing interest.

